# Discovering and Validating Neoantigens by Mass Spectrometry-based Immunopeptidomics and Deep Learning

**DOI:** 10.1101/2022.07.05.497667

**Authors:** Ngoc Hieu Tran, Chao Peng, Qingyang Lei, Lei Xin, Jingxiang Lang, Qing Zhang, Wenting Li, Haofei Miao, Ping Wu, Rui Qiao, Haiming Qin, Dongbo Bu, Haicang Zhang, Chungong Yu, Xiaolong Liu, Yi Zhang, Baozhen Shan, Ming Li

## Abstract

Neoantigens are promising targets for cancer immunotherapy but their discovery remains challenging, mainly due to the sensitivity of current technologies to detect them and the specificity of our immune system to recognize them. In this study, we addressed both of those problems and proposed a new approach for neoantigen identification and validation from mass spectrometry (MS) based immunopeptidomics. In particular, we developed DeepNovo Peptidome, a *de novo* sequencing-based search engine that was optimized for HLA peptide identification, especially non-canonical HLA peptides. We also developed DeepSelf, a personalized model for immunogenicity prediction based on the central tolerance of T cells, which could be used to select candidate neoantigens from non-canonical HLA peptides. Both tools were built on deep learning models that were trained specifically for HLA peptides and for the immunopeptidome of each individual patient. To demonstrate their applications, we presented a new MS-based immunopeptidomics study of native tumor tissues from five patients with cervical cancer. We applied DeepNovo Peptidome and DeepSelf to identify and prioritize candidate neoantigens, and then performed *in vitro* validation of autologous neoantigen-specific T cell responses to confirm our results. Our MS-based *de novo* sequencing approach does not depend on prior knowledge of genome, transcriptome, or proteome information. Thus, it provides an unbiased solution to discover neoantigens from any sources.

## Introduction

Tumor-specific human leukocyte antigen (HLA) peptides are only expressed on the surface of cancer cells and hence represent ideal targets for the immune system to fight cancer. However, due to the high specificity of T cells, only a very small fraction of those peptides can be recognized by T cells to trigger immune responses. They are referred to as neoantigens. Three landmark studies by Carreno et al.^1^, Ott et al.^2^ and Sahin et al.^3^ have successfully developed personal neoantigen vaccines in clinical trials with melanoma patients. In those studies, DNA and RNA sequencing were first used to identify somatic mutations. Subsequently, *in silico* methods^4,5^ were used to predict which mutated peptides were more likely to bind to the HLA proteins and to be recognized by the T cells of the patient. Up to 20 candidate neoantigens per patient were finally selected to manufacture the personal vaccine. In another milestone, Bassani-Sternberg et al.^6^ reported the first identification of neoantigens directly from native tumor tissues by mass spectrometry (MS) based immunopeptidomics. The study provided direct evidence of neoantigen presentation rather than just *in silico* predictions. Since then, MS-based immunopeptidomics has increasingly become the method of choice for neoantigen discovery.

Despite their promising role in cancer immunotherapy, the chance of finding neoantigens is remarkably low, usually less than half a dozen out of thousands of somatic mutations detected per patient^7–10^. There are two major challenges that we need to address: sensitivity and specificity. The first challenge is the sensitivity of MS-based immunopeptidomics. Tumor-specific HLA peptides may arise from somatic mutations in coding regions, or from non-coding regions such as lncRNAs, transposable elements (TEs), unannotated open reading frames (ORFs), etc. Hence, a common approach in proteogenomics studies is to integrate multi-omics data, including genomics, transcriptomics, and ribosome profiling, to build large customized protein databases to search against MS data. Nevertheless, very few non-canonical HLA peptides could be detected^6,11–13^, even when the false discovery rate (FDR) was relaxed to 3-5% instead of the typical 1%. It is still unclear how to address that limitation, and current solutions such as building larger databases or increasing the FDR seem not good enough. On the other hand, recent studies have found that many HLA peptides exhibited minimal RNA expression^14,15^. Those observations suggest that a different approach independent from transcriptomics, such as *de novo* peptide sequencing, is highly desired to identify non-canonical HLA peptides.

The second challenge is the specificity of T cells. Majority of current *in silico* methods focus on predicting the binding affinity or the presentation of class 1 and class 2 peptide-HLA (pHLA) complexes^4,16–20^. However, such approaches only consider half of the equation and do not take into account how a pHLA complex can be recognized by T cell receptors (TCRs). Previous studies on HLA class 1 (HLA-I) T cell epitopes have suggested that certain amino acid positions, e.g. 4-6, of the epitopes were in close contact with the TCRs, while the anchor positions, e.g. 1, 2, 9, were responsible for HLA-I binding^21–24^. In addition, certain properties of amino acid residues, such as hydrophobicity, polarity, or large and aromatic side chains, were found to have statistically significant correlation with the epitope immunogenicity. Sequence similarity to known pathogen epitopes and dissimilarity to the human proteome have also been proposed for immunogenicity prediction^25,26^. However, those methods still fall short of accounting for the specificity of T cells in each individual patient.

In this study, we aim to address both of the aforementioned sensitivity and specificity challenges. First, we proposed DeepNovo Peptidome, a *de novo* sequencing-based search engine for HLA peptide identification. Our MS-based *de novo* sequencing approach is independent from genome, transcriptome, or proteome databases. Thus, it could identify any non-canonical HLA peptides regardless of their sources. To boost the sensitivity of HLA peptide identification, we collected a massive dataset of 26 millions MS/MS spectra and 1.1 millions unique HLA peptides to train a deep learning-based de novo sequencing model^27,28^. The model can learn the patterns of HLA binding motifs to enhance its de novo sequencing capability.

Those motifs are especially critical for *de novo* sequencing of HLA peptides because they are located at the second amino acid position, which often lacks supporting *b/y* fragment ions in MS/MS spectra^15^. Once non-canonical HLA peptides have been identified by DeepNovo Peptidome, it is essential to predict their immunogenicity and to prioritize top candidates for *in vitro* validations. We proposed DeepSelf, a personalized model for immunogenicity prediction based on the central tolerance of T cells, i.e. the positive and negative selection of T cells in an individual patient. In the positive selection, T cells are selected by their ability to bind to peptide-HLA complexes. In the negative selection, they are selected against their ability to bind to self peptides. Thus, the key idea of DeepSelf is to learn HLA self peptides obtained from MS-based immunopeptidomics of each individual patient to resemble the negative selection of the T cells of that patient.

DeepNovo Peptidome and DeepSelf together form a complete solution for identifying non-canonical HLA peptides and predicting their immunogenicity. We demonstrated the high sensitivity of DeepNovo Peptidome by applying it to the HLA Ligand Atlas^29^, a public immunopeptidomics database of benign human tissues. We identified 181,012 HLA-I and 208,255 HLA-II peptides, substantially expanding the HLA Ligand Atlas by 46-100%. We also extensively evaluated DeepSelf on the immunopeptidomes and neoantigens of 18 cancer patients from 10 previously published studies^2,6,16,30–36^. Finally, we presented a new MS-based immunopeptidomics study of native tumor tissues from five patients with cervical cancer. We performed a comprehensive experiment and analysis, from sample preparation to peptide identification, tumor antigen profiling, and immunogenicity prediction. We prioritized candidate neoantigens from the tumor samples, performed *in vitro* validations, and confirmed four neoantigens with autologous neoantigen-specific T cell responses.

## Results

### Boosting the sensitivity of MS-based immunopeptidomics with DeepNovo Peptidome

Since neoantigens arise from non-canonical sources which are challenging for standard database search, we built DeepNovo Peptidome, a *de novo* sequencing-based search engine for HLA peptide identification and neoantigen discovery. We trained a deep learning-based *de novo* sequencing model^27,28^ on a very large, carefully curated dataset from 20 previous MS-based immunopeptidomics studies^6,17–19,29,30,37–50^. The dataset included nearly 3,000 runs on two MS instruments Orbitrap and timsTOF, >26M spectra and >1.1M unique HLA peptides, and covering more than 90 alleles for each HLA class. The newly trained model, named DeepNovo-HLA, could learn the patterns of fragment ions and binding motifs of HLA peptides from the training data. It then used that knowledge to perform *de novo* sequencing of non-canonical HLA peptides. In addition, database search and spectral library search^20^ were also integrated into DeepNovo Peptidome to identify canonical HLA peptides. Finally, canonical and non-canonical HLA peptides were combined and evaluated by a joint FDR estimation to produce a complete immunopeptidome. A summary of DeepNovo Peptidome search engine is depicted in Supplementary Figure S1.

### Expanding the HLA Ligand Atlas of benign human tissues by two-fold

We next demonstrated the application of DeepNovo Peptidome to expand the HLA Ligand Atlas, a database of HLA-I and HLA-II immunopeptidomes from >200 benign human tissue samples published recently by Marcu et al^29^. Such a benign database is essential as it can be used as a reference to compare to the immunopeptidomes obtained from tumor tissues. The reference benign database can help to improve the safety of T-cell-based immunotherapies by avoiding potential on-target off-tumor adverse events. It can also be used to study the differences between possible pairs of wild-type and mutated HLA peptides and how such differences influence their binding and immunogenicity^51^. The current HLA Ligand Atlas includes 90,428 HLA-I and 142,625 HLA-II peptides from 51 HLA-I and 86 HLA-II allotypes (https://hla-ligand-atlas.org).

Given the critical role of the HLA Ligand Atlas for immunotherapies, here we leveraged the high sensitivity of DeepNovo Peptidome to substantially expand this reference database. We downloaded >1200 LC-MS/MS runs from that study and performed DeepNovo Peptidome search with precursor tolerance 10 ppm,fragment ion tolerance 0.02 Da, variable Met(Oxidation), peptide length 8-14 for HLA-I and 8-25 for HLA-II, and peptide FDR at 1%. The search parameters were set similar to those used in the original study. The search results are summarized in Figure 1. DeepNovo Peptidome identified 181,012 HLA-I and 208,255 HLA-II peptides, substantially expanding the HLA Ligand Atlas. The number of HLA-I peptides was increased by two folds and the number of HLA-II peptides was increased by 46% (Venn diagrams in Figure 1a, d). Our results also expanded the Immune Epitope Database^5^ (IEDB) by nearly 73K HLA-I peptides and 60K HLA-II peptides. We also validated the length distributions and the binding affinities of the identified peptides to their respective HLA alleles using NetMHCpan and NetMHCIIpan^4^. It can be seen that the numbers of HLA peptides were consistently increased across different lengths and different alleles of both HLA classes, demonstrating the sensitivity and robustness of DeepNovo Peptidome (Figure 1b, c, e, f). The details of the identified peptides and their peptide-spectrum matches (PSMs) are provided in Supplementary Data 1.

**Fig. 1.**
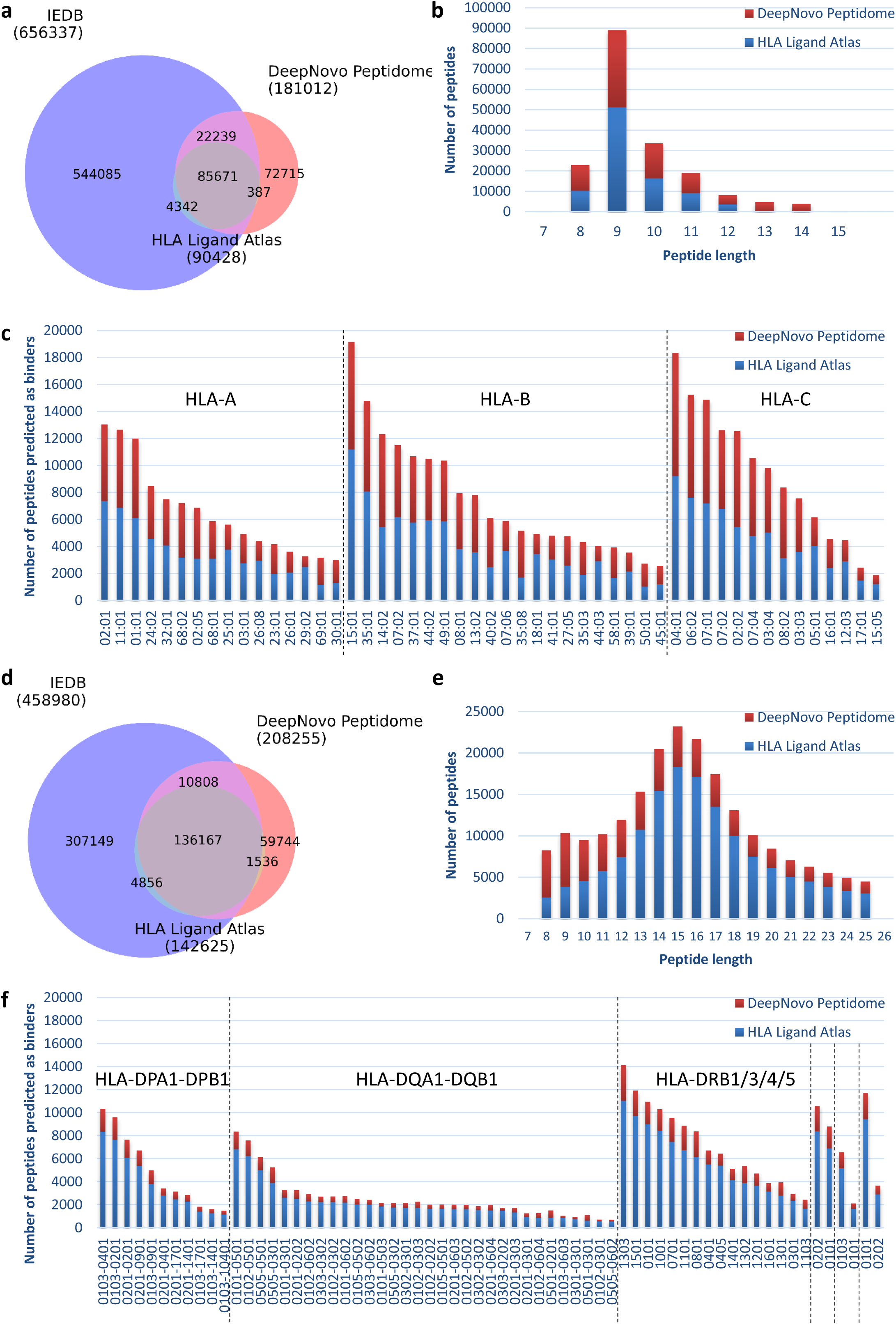
DeepNovo Peptidome workflow substantially expanded the reference HLA Ligand Atlas, increasing the number of HLA-I peptides by two folds and the number of HLA-II peptides by 46%. **a-c**, Venn diagram, length distribution, and NetMHCpan predicted binders of HLA-I peptides. d-f, Venn diagram, length distribution, and NetMHCIIpan predicted binders of HLA-II peptides. (HLA: Human Leukocyte Antigen; IEDB: Immune Epitope Database)

### DeepSelf: a personalized model for immunogenicity prediction based on the central tolerance of T cells

Once the immunopeptidome of a patient had been identified by DeepNovo Peptidome, the non-canonical HLA peptides could be considered as candidate neoantigens of that patient. We further used DeepSelf to predict their immunogenicity and prioritize top candidates for *in vitro* validations. DeepSelf is a personalized model for immunogenicity prediction based on the central tolerance, i.e. the positive and negative selection of T cells in an individual patient (Figure 2). In the positive selection, T cells are selected by their ability to bind to peptide-HLA complexes. In the negative selection, they are selected against their ability to bind to self peptides. Thus, we proposed to use HLA self peptides obtained from MS-based immunopeptidomics to resemble the negative selection of T cells in each individual patient. For the positive selection, we collected all positive T cell epitopes from the IEDB that matched the patient’s HLA alleles. DeepSelf personalized model was then trained on those negative self peptides and positive T cell epitopes of the patient to predict the immunogenicity of his/her candidate neoantigens. Our model did not impose pre-defined features of amino acid properties or positions like previous immunogenicity prediction tools^5,23,24^. Instead, we used a bi-directional LSTM network coupled with amino acid embedding to model the negative and positive peptides. This approach had been successfully used previously for de novo sequencing^27,52^, spectrum and retention time predictions^37,53,54^, and other peptide prediction tasks^8,55^. More details of the DeepSelf model and training can be found in the Methods section.

**Fig. 2.**
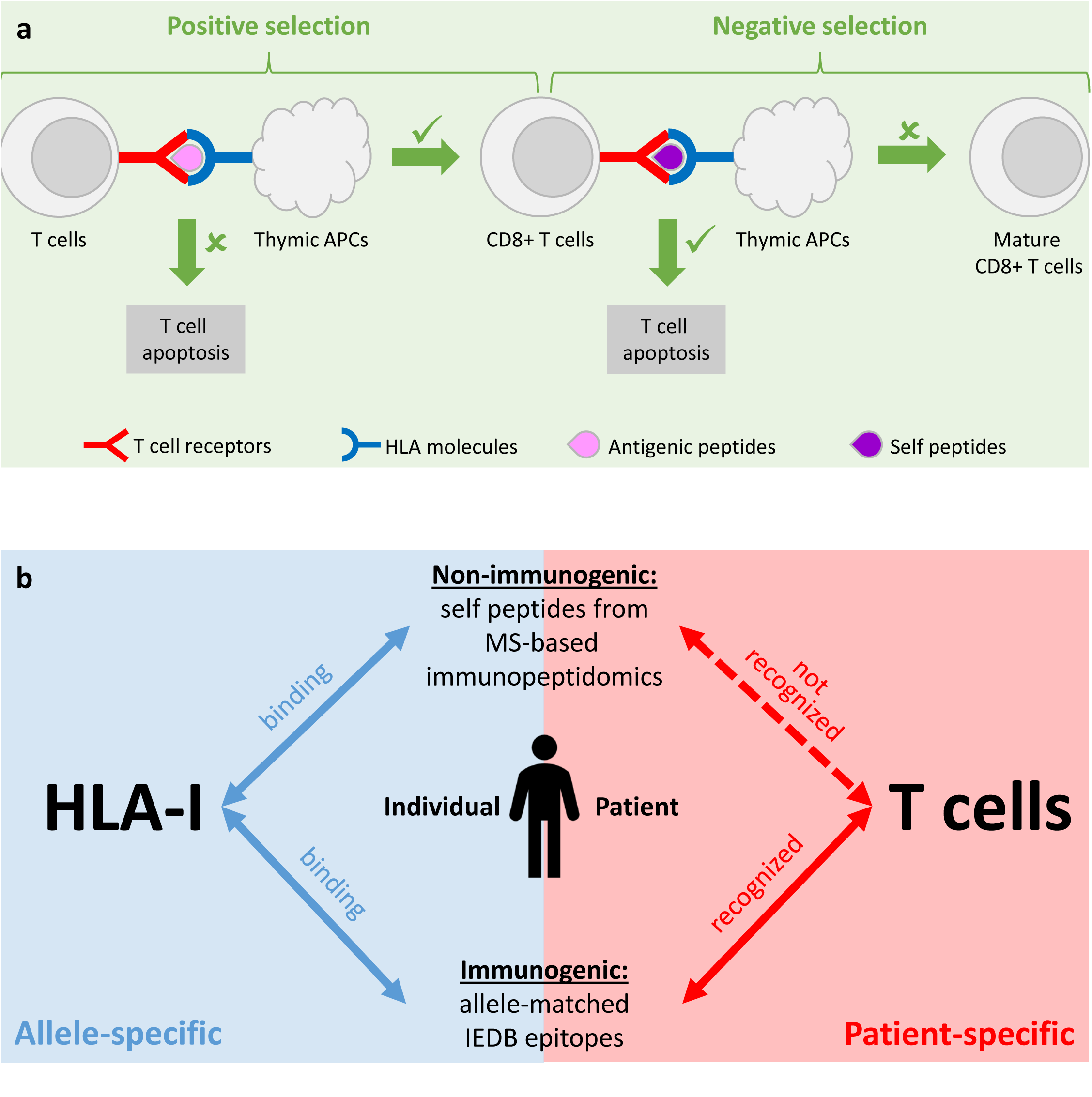
DeepSelf, a personalized model for immunogenicity prediction based on the central tolerance of T cells. **a**, The positive and negative selection processes, i.e. the central tolerance of CD8+ T cells. During the positive selection, T cells are selected by their ability to bind to peptide-HLA complexes: they will become CD8+ T cells if they bind to HLA-I complexes, otherwise they will die by apoptosis. During the negative selection, CD8+ T cells are further selected *against* their ability to bind to self peptides: those that have high affinity for self peptides will die by apoptosis, preventing the risk of autoimmunity; the remaining will become mature CD8+ T cells. **b**, Personalized immunogenicity prediction model to resemble the central tolerance of CD8+ T cells in an individual patient. We use MS-based immunopeptidomics to obtain HLA-I self peptides to resemble the negative selection of CD8+ T cells in an individual patient. For the positive selection, we collect all epitopes reported in positive T cell assays from IEDB that match the patient’s HLA-I alleles. Using this personal dataset of positive and negative peptides, we then train a binary classification model specifically for that patient to predict his/her T cell response to any given peptide. (MS: Mass Spectrometry; HLA: Human Leukocyte Antigen; APC: Antigen-Presenting Cell; IEDB: Immune Epitope Database)

DeepSelf was rigorously evaluated on the immunopeptidomes and neoantigens of 18 cancer patients from 10 previously published studies^2,6,16,30–36^ (Methods, Figure 3). The number of neoantigens varied from 2-6 per patient, all of which had been confirmed as immunogenic by T cell assay validation (Supplementary Table S3). We also compared DeepSelf to three other popular tools for immunogenicity prediction, including PRIME^23,24^ (version 2.0), NetMHCpan^4^ (version 4.1), and IEDB immunogenicity predictor^21^. DeepSelf and three other prediction tools were tasked to prioritize 1 immunogenic neoantigen out of every 100 non-immunogenic HLA peptides. DeepSelf achieved an average accuracy of 0.69 and outperformed the other tools on 12 out of 18 patients (Figure 3a, c, Supplementary Data 3). It was able to rank 25% of the neoantigens within its top 10% predictions, and nearly 75% of the neoantigens within its top 30% (Figure 3b). In other words, for an average patient evaluation set that included 4 neoantigens and 400 non-immunogenic peptides, DeepSelf’s top 40-120 predicted candidates would contain 1-3 neoantigens of interest. In a special case of patient Mel-15 with the deepest immunopeptidome coverage, DeepSelf was able to rank 3 neoantigens within its top 3% predictions (Figure 3d). More details of the evaluation datasets and criteria can be found in the Methods section.

**Fig. 3.**
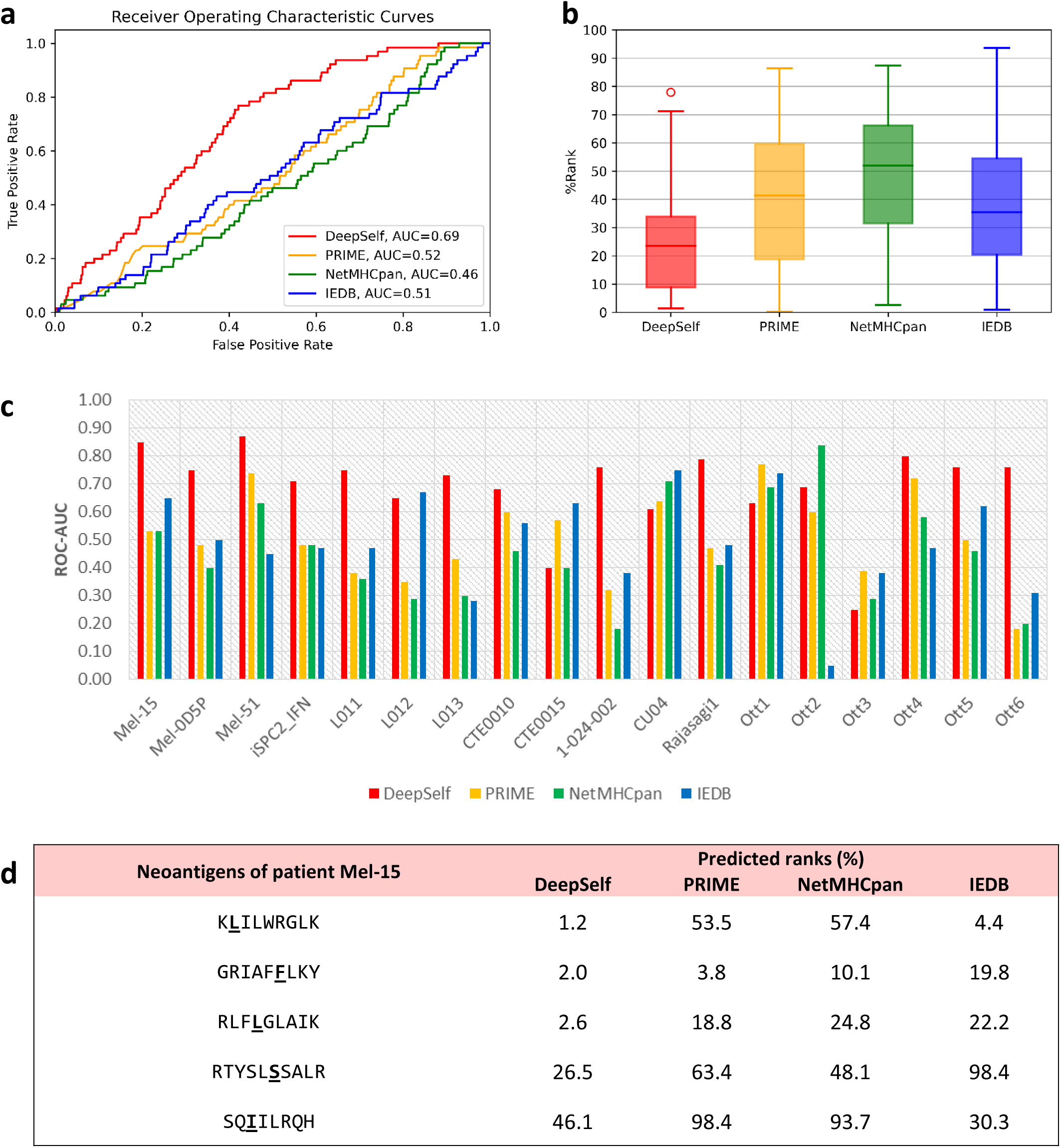
Performance evaluation of four immunogenicity prediction tools on 18 cancer patients. **a,** Receiver operating characteristic curves on the combined evaluation set of 18 patients. **b,** Predicted ranks of immunogenic neoantigens versus non-immunogenic peptides (lower % is better). **c,** Areas under the receiver operating characteristic curves (ROC-AUC) of the prediction tools on each individual patient. **d,** An example of predicted ranks of patient Mel-15’s neoantigens. (IEDB refers to the immunogenicity prediction tool on IEDB website developed by Calis et al., 2013. Bold underlined letters in panel d indicate mutated amino acids).

### Identifying and validating neoantigens from native tumor tissues of five patients with cervical cancer

In the next sections, we present a new MS-based immunopeptidomics study of native tumor tissues from five patients with cervical cancer. The patient information is summarized in Table S1. The patients were enrolled under the protocols approved by the Ethics Committees at the First Affiliated Hospital of Zhengzhou University, China. We demonstrated how DeepNovo Peptidome and DeepSelf could be applied to identify and validate neoantigens from these samples.

### MS-based immunopeptidomics analysis of native tumor tissues

Frozen cervical cancer tissues from five patients were selected for neoantigen discovery. The amount of the tissues varied from 18mg to 110mg. An immunopeptidome enrichment kit, named Neo Discovery, was developed and used for the sample preparation. Briefly speaking, the biopsy or surgical cutting block tissues were first homogenized to extract HLA-peptide complexes with the lysis buffer in the kit following the manufacturer instructions. To ensure the stability of HLA-peptide complexes and thorough extraction of membrane protein complexes from the tissues, the samples were kept at 4 degree during the whole process and different combinations of multiple detergents were optimized for the complete membrane complex solubilization and extraction. Different kinds of pan-antibodies for HLA-peptide complex recognition and enrichment were also developed and evaluated to increase the yield of HLA-peptide complexes. In addition, crosslinking strategies for antibodies and beads, such as different types of linkers and beads were also tested to increase the crosslinking efficiency and the activity of antibodies after the crosslinking. The HLA-peptide complexes were eluted from the beads by acid solutions and then cleaned up with a reverse phase column. The immunopeptides were eluted by low concentration of acetonitrile solution and HLA proteins were eluted by high concentration of acetonitrile solution. 5% of the samples of each fraction from each step were collected for quality control. Based on our Western Blot validation results, above 90% of the HLA-peptide complexes were extracted in the supernatant of the tissue lysate and above 95% of the complexes were enriched.

The enriched immunopeptides were dried out for subsequent LC-MS/MS analysis. TimsTOF Pro2 mass spectrometer with PASEF mode was used for the detection of low-abundance immunopeptides with high sensitivity. Specific parameters were also optimized for the timsTOF Pro2 instrument for the analysis of immunopeptides. The acquired MS data of each patient was further analyzed using DeepNovo Peptidome and DeepSelf for neoantigen identification and immunogenicity prediction (Methods).

We identified a total of 58,756 unique HLA-I peptides, where the number of peptides per patient varied from 7,682-22,220, mainly due to the available amount of tumor tissues (Figures 4a, b). 73% of the peptides were found only in one patient, 21% were shared by two patients and 6% were shared by 3-5 patients. Figure 4c shows that the peptides identified from all five patients followed the HLA-I characteristic length distribution with majority of them being 9-mers. 91-95% of the peptides could be found in the human protein database (i.e. canonical HLA peptides), whereas the remaining 5-9% were *de novo* peptides (i.e. non-canonical HLA peptides) (Figure 4d). We also checked the patient peptidomes against the IEDB and found that 44-56% of the peptides had been reported in the IEDB, and the remaining 43-56% were new epitopes (Figure 4e). Notably, we found 18-71 peptides (<0.5%) that had been reported as positive T cell epitopes in previous studies. The relative ratios between T cell epitopes and IEDB epitopes were extremely low, less than 1:100, reiterating the challenge of finding immunogenic neoantigens.We also used NetMHCpan^4^ to predict the binding of the identified peptides to the HLA molecules of each patient. Figure 4f shows that 63-90% of the peptides were predicted as binding (%rank < 2), and among them, 56-81% of the peptides were predicted as strong binding (%rank < 0.5). Overall, the results presented in Figure 4 demonstrate the high sensitivity and accuracy of DeepNovo Peptidome on real clinical samples of native tumor tissues from cervical cancer patients.

**Fig. 4.**
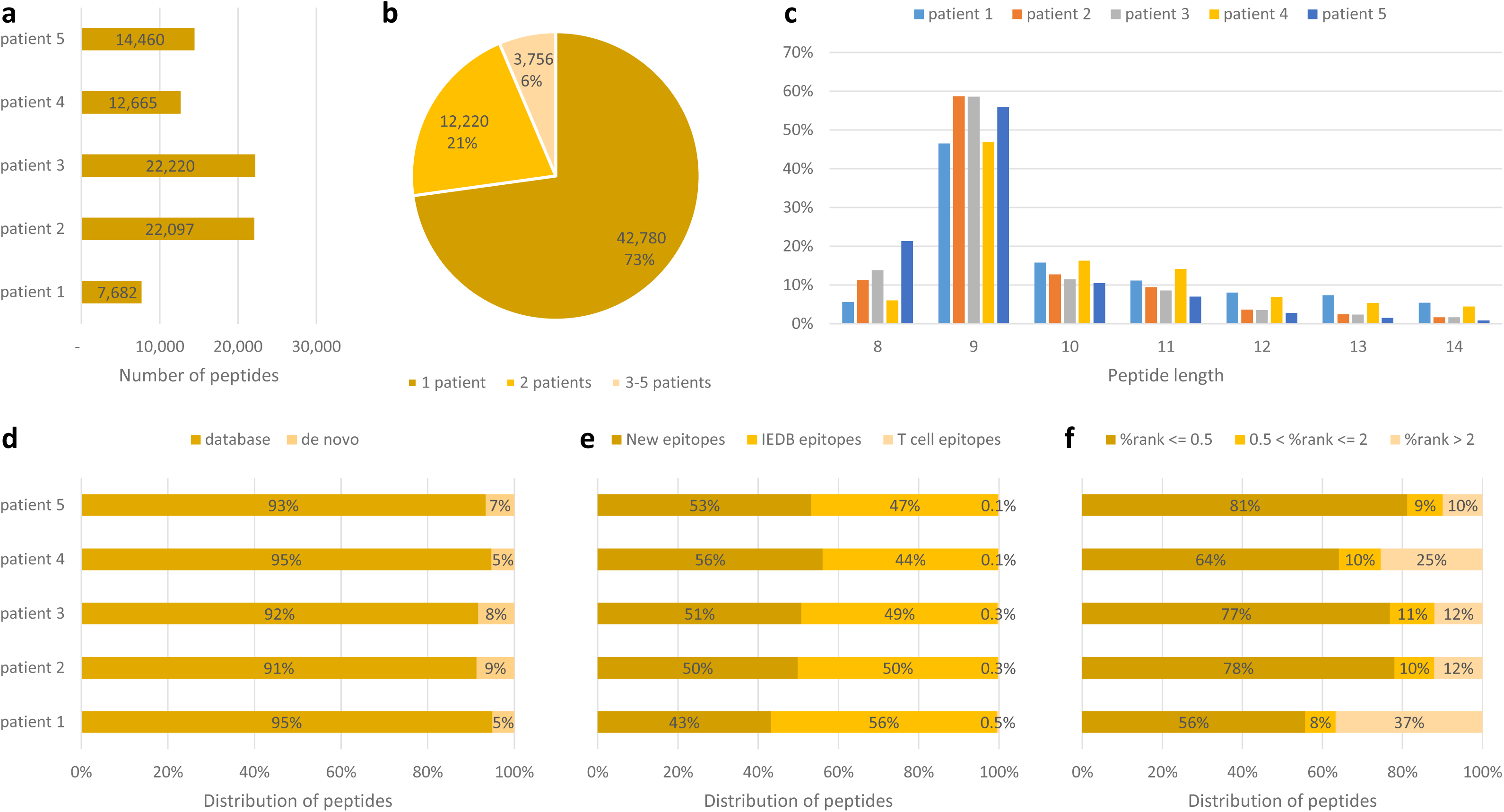
Summary of the HLA-I peptidomes identified from MS-based immunopeptidomics analysis of native tumor tissues of 5 patients with cervical cancer. **a,** Number of identified HLA-I peptides per patient. **b,** Unique and shared HLA-I peptides in the cohort. **c,** Length distribution of HLA-I peptides. **d**, Proportions of database and de novo HLA-I peptides. **e**, Proportions of known IEDB epitopes and positive T cell epitopes among the identified HLA-I peptides. The remaining HLA-I peptides are considered as new epitopes. **f**, Predicted binding %rank of HLA-I peptides by NetMHCpan. (MS: Mass Spectrometry; HLA: Human Leukocyte Antigen; IEDB: Immune Epitope Database)

### Presentation of HLA peptides from tumor-associated antigens

While the main focus of our study is neoantigens, HLA peptides that are derived from tumor-associated antigens (TAAs) and are shared between multiple patients may also represent potential targets for cancer immunotherapy. We collected a list of 940 TAAs from previous cancer immunology studies^6,11,12^ and analyzed their HLA peptide presentation in the five cervical cancer patients in our study (Figure 5). We found a total of 473 unique HLA-I peptides that were derived from 130 TAAs in the five patients. While the majority of the peptides were unique to one patient, a large number of TAAs were shared between multiple patients (Figures 5b, c).

**Fig. 5.**
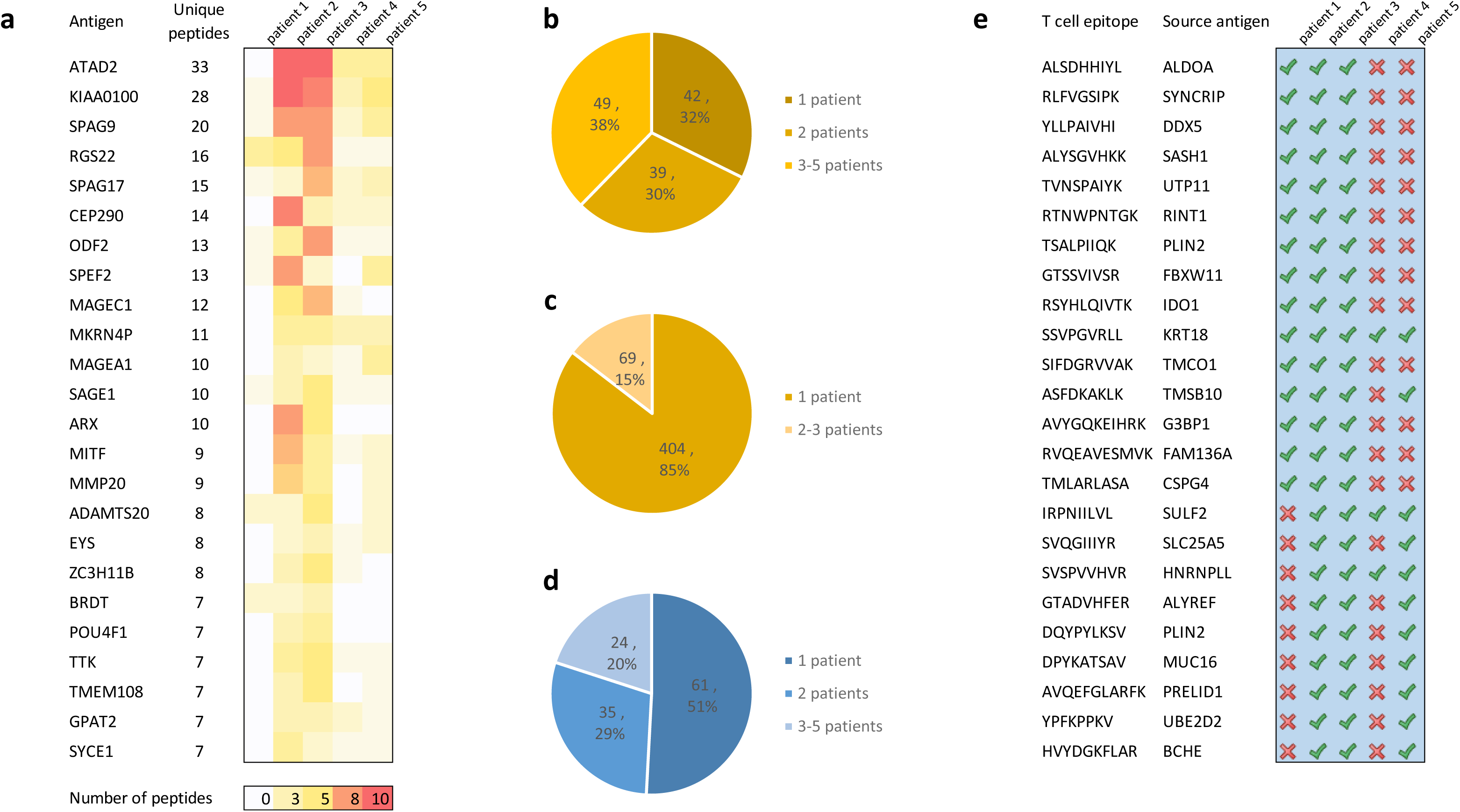
Presentation of TAA-derived HLA-I peptides and known T cell epitopes that were found in the tumor tissues of 5 cervical cancer patients. **a,** Heat map of the number of HLA-I peptides per patient that were derived from top 24 TAAs. **b,** Distribution of unique and shared TAAs in the cohort. **c,** Distribution of unique and shared TAA-derived HLA-I peptides in the cohort. **d**, Distribution of unique and shared T cell epitopes in the cohort. **e**, Common T cell epitopes that were found in the majority of the patients. (TAA: Tumor-Associated Antigen)

Figure 5a shows the heat map of HLA-I peptide presentation of the top 24 TAAs with the highest number of unique peptides. The top two genes, *ATAD2* and *BLTP2* (KIAA0100) are well-known for their association with endometrial and breast cancers. In addition to TAA-derived peptides, we also investigated the HLA-I peptides that had been reported as positive T cell epitopes by previous studies in the IEDB. We found a total of 120 positive T cell epitopes in the patients’ peptidomes, and more interestingly, 24 of them were shared by 3-5 patients (Figures 5d, e). The presentation of those common TAAs and T cell epitopes suggests that they may represent promising candidates for off-the-shelf immunotherapy treatments, which may be effective across a number of different patients, as opposed to neoantigens that are unique to an individual patient.

### Immunogenicity prediction and *in vitro* validation of autologous neoantigen-specific T cell responses

Once the HLA-I peptidome had been identified by DeepNovo Peptidome for each cervical cancer patient, the non-canonical HLA peptides were further considered as candidate neoantigens, whereas the canonical HLA peptides were considered as self peptides (Figure 4d). Positive training data (T cell epitopes) for each patient were obtained from the IEDB according to the patient’s alleles. The number of self peptides varied from 7,293-20,367 and the number of positive T cell epitopes varied from 536-4,407 (Table S1). Personalized DeepSelf models were then trained for each individual patient and subsequently applied to predict the immunogenicity of his/her candidate neoantigens. The candidate neoantigens and prediction results are provided in the Supplementary Data 2.

To further validate our neoantigen prediction results, we performed *in vitro* validation of autologous neoantigen-specific T cell responses on the samples of patient 1. This patient had been diagnosed with stage-III cervical cancer and had received chemotherapy, radiotherapy and PD-1 therapy. The details of her clinical information and biomaterial sample collection are shown in Figure 6a. The tissue sample collected before the first cycle of radio-immunotherapy was used to identify the candidate neoantigens. As summarized in Table S1, 7,682 HLA-I peptides were identified by DeepNovo Peptidome from this patient. Among those, 7,293 canonical peptides were considered as self peptides and 389 non-canonical peptides were considered as candidate neoantigens. The patient’s self peptides and 4,266 allele-matched T cell epitopes from the IEDB were used to train the DeepSelf personalized model for this patient. The model was then used to rank the 389 non-canonical peptides. In addition to the default ranking by DeepSelf, we also tried to balance the number of positive T cell epitopes due to the over-representation of the patient’s allele HLA-A*02:01 in the IEDB. The top 100 non-canonical peptides were finally synthesized for autologous TCR stimulation tests. The peripheral blood mononuclear cells (PBMC) were obtained from the patient blood sample as described in Figure 6b. Four peptides were detected as positive with IFN-γ ELISpot multiple times, with significantly more spots compared to negative controls (Figure 6c). They ranked at the 38, 39, 81, and 82 places among the candidate neoantigens of this patient (Table S2). We also assessed the expression of CD137 on the patient’s autologous T cells after stimulation with these four peptides, as well as the expression of cytokines on the CD137+ T cells by flow cytometry. We found that the proportion of neoantigen-specific T cells was increased after stimulation and that there was also a significant increase in the expression of IFN-γ (Figure 6d-f).

**Fig. 6.**
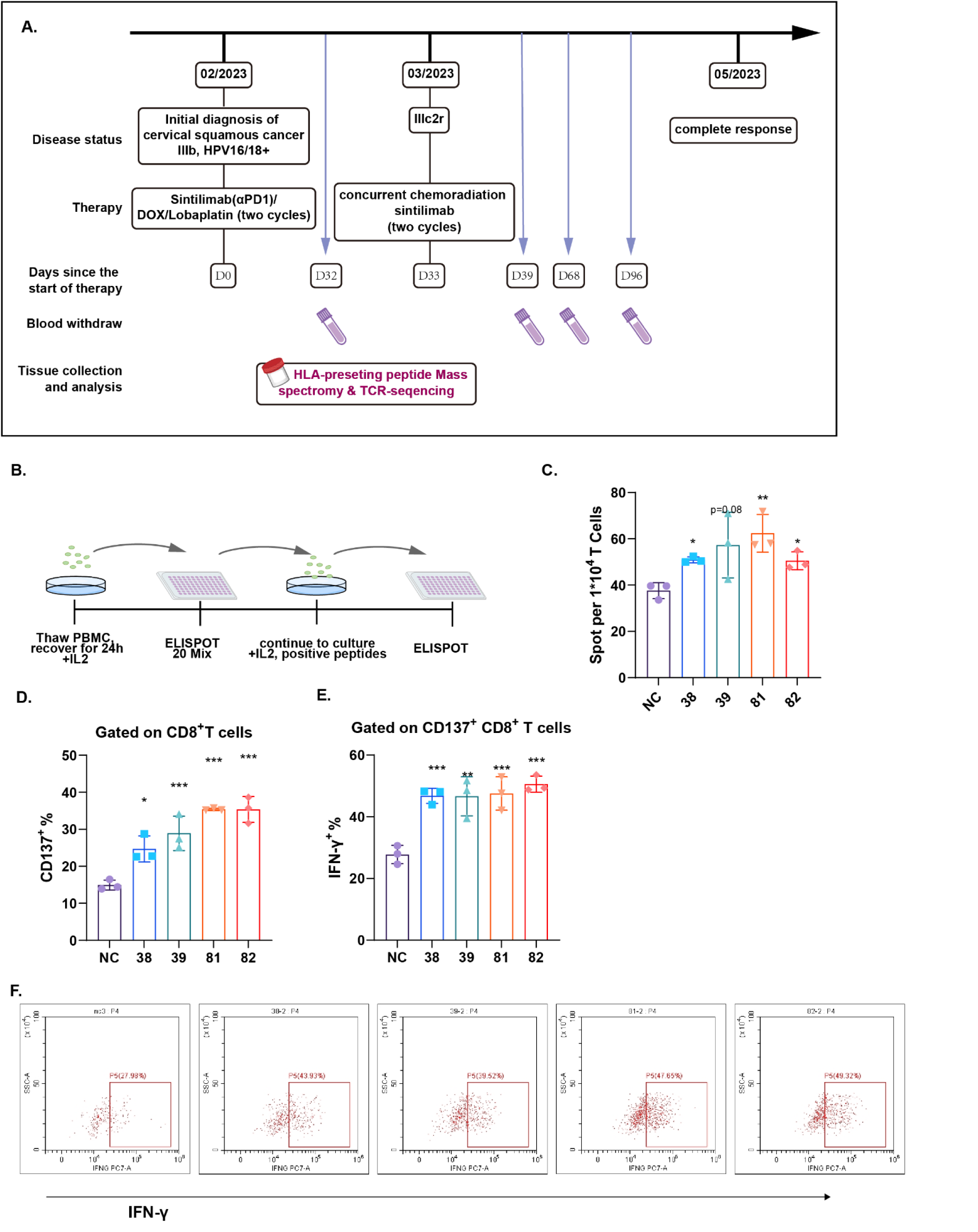
*In vitro* validation of autologous neoantigen-specific T cell responses on the samples of patient 1 with cervical cancer. **a**, Clinical information of the patient. **b**, Overview of the ELISpot the experiments with PBMC. **c,** Spots formation of blood-derived PBMC against the four identified neoantigens from patient 1, adjusted to 10^4^ cells. **d,** CD137 expression of epitope mixture expanded PBMC after incubation with four identified neoantigens (mean ± s.e.m. of 3 replicates). **e,** Intracellular cytokine staining (ICS) of expanded PBMC after epitopes exposure gated on CD137+ T cells (mean ± s.e.m. of 3 replicates). **f,** Representative image of flow cytometry detecting IFN-γ on neoantigen-specific T cells (mean ± s.e.m. of 3 replicates). **d-f,** D’Agostino-Pearson omnibus normality test was conducted to confirm the distribution of the data, and all the data passed the normality test. P-value was determined with a two-tailed unpaired t-test. *: P<0.05, **: P<0.01; ***: P<0.001.

## Discussion

In this study, we proposed DeepNovo Peptidome and DeepSelf, which together form a complete solution for discovering and validating neoantigens from MS-based immunopeptidomics. Our *de novo* sequencing-based approach can identify neoantigens directly from native tumor tissues and does not depend on genome or transcriptome information. Thus, it represents a promising approach to overcome current limitations of proteogenomics studies, such as limited sensitivity of detecting non-canonical HLA peptides, and/or minimal expression of many HLA peptides at the transcriptome level^9,14,15^.

We demonstrated that, for a given sample of tumor tissue, DeepNovo Peptidome and DeepSelf were able to identify and prioritize 100 candidate neoantigens for *in vitro* T cell validation, and four of them were confirmed to be immunogenic. At the time of submitting this manuscript, an independent study by Gurung et al.^56^ has also reported a successful application of DeepNovo Peptidome for systematic discovery of neoepitope–HLA pairs for neoantigens shared among patients and tumor types. The results together confirm the validity and the advantage of our approach for neoantigen discovery in cancer immunotherapy.

While the data and analysis in this study mainly focused on cancer patients, we believe that our MS-based approach for neoantigen discovery also offers a promising resource for the development of treatments for autoimmune diseases. A significant number of autoimmune diseases are caused by HLA abnormalities and hence standard HLA binding prediction tools may no longer provide accurate results. In such cases, MS-based immunopeptidomics is the only way to interrogate the patient’s immunopeptidome and to identify the neoantigens causing the diseases^57–60^.

Our immunogenicity prediction tool DeepSelf considers the patient’s self peptides identified from MS as non-immunogenic (negative training data), and immunogenic T cell epitopes from the IEDB as positive training data. For patients lacking MS data and self peptides, HLA allele-matched peptides from the IEDB can be used as negative training data. Using self peptides did indeed provide more accurate predictions than using HLA allele-matched peptides, as we have demonstrated in comparison to other allele-based tools like NetMHCpan and PRIME. We have also tested our own model using both types of negative training data and found that using self peptides yielded better results.

As DeepSelf selects immunogenic T cell epitopes from the IEDB based on the six alleles of a patient, a possible limitation is that, for some uncommon alleles, there are relatively few immunogenic epitopes. In addition, while we have tried to search for as many patient datasets as possible for evaluation, the results in this study were based on the datasets of 18 cancer patients from 10 previously published studies^2,6,16,30–36^. The number of neoantigens varied from 2-6 per patient, and only four patients had MS-based immunopeptidomics data available. Thus, those limitations need to be acknowledged when interpreting the evaluation results in this study.

## Methods

### Tumor tissues and blood samples

After 2 cycles of chemotherapy, tumor tissues were taken with biopsy forceps and weighed with an analytical balance, prior to radiotherapy. Tissues were immediately cut into 2 pieces for TCR sequencing (67mg) and antigen detection (18mg), respectively. Blood samples were taken before, during and after radiotherapy. Peripheral blood mononuclear cells (PBMCs) were isolated from blood samples immediately after collection and cryopreserved in 90% bovine serum + 10% DMSO.

### HLA peptide purification and LC-MS/MS analysis

MHC-peptide complexes were enriched with our Neo Discovery^TM^ immunopeptidome enrichment Kit (BaizhenBio Inc.). The tissue samples were homogenized in a grinding mill with liquid nitrogen. Lysis buffer was then added into the pellet and incubated on ice for 1 hour. The mixtures were sonicated for 3 minutes on ice and centrifuged at 20000g, 4℃ for 1 hour. The supernatant was aspirated for MHC-peptide complex enrichment with anti-HLA I antibody crosslinked beads included in the kit. The immuno-peptides were then separated and desalted by a C18 spin column (GL science). Eluted fractions containing immunopeptide were dried out by a speed-vacuum instrument.

Peptides were reconstituted by 0.1% Formic Acid, and separated on a homemade C18 column (200 mm x 75 μm i.d., 1.9 Å particle size; Dr. Maisch GmbH) by a nanoElute HPLC (Bruker) at 300nL/min with a 60 min gradient of 4-100% Acetonitrile (with 0.1% formic acid). Data were acquired by a timsTOF Pro2 Mass Spectrometer (Bruker) in a data-dependent method with 10 PASEF (Parallel Accumulation–Serial Fragmentation). Mass Range was set as 300-1700m/z, ion threshold for MS2 was set as 2500. The acquired MS data of each patient was further analyzed using DeepNovo Peptidome and DeepSelf for neoantigen identification and immunogenicity prediction, as described below. The precursor tolerance and fragment ion tolerance were set at 20 ppm and 0.05 Da, respectively; Met(Oxidation) was set at variable modification.

### Training personalized DeepSelf models for immunogenicity prediction

The specificity of T cells is mainly shaped by their positive and negative selection, i.e. the central tolerance that happens inside the thymus of each individual patient (Figure 2a). During the positive selection, T cells are selected by their ability to bind to peptide-HLA complexes: they will become CD8+ or CD4+ T cells if they bind to HLA-I or HLA-II complexes, respectively; otherwise they will die by neglect^61^. During the negative selection, T cells are selected against their ability to bind to self peptides: those that have high affinity for self peptides will die by clonal deletion, preventing the risk of autoimmunity; the remaining will mature and participate in immune responses against foreign antigens. Motivated by the central tolerance of T cells, we hypothesize that the positive and negative selection can be encapsulated by the complete space of positive and negative peptides resulting from that selection. Thus, we propose to use MS-based immunopeptidomics to obtain HLA self peptides to resemble the negative selection of T cells in an individual patient (Figure 2b). For the positive selection, one can collect all epitopes reported in positive T cell assays from the Immune Epitope Database (IEDB)^5^ that match the patient’s HLA alleles. Both self peptides and T cell epitopes bind to the HLA molecules of the patient, however, the former are non-immunogenic and the latter are immunogenic to the patient’s T cells. Using this personal dataset of negative and positive peptides, a binary classification model can be trained specifically for that patient to predict his/her T cell responses to any given peptide.

In particular, for each individual patient in this study, the MS-based immunopeptidomics data was analyzed using DeepNovo Peptidome described in the previous sections. The identified peptides were further checked for their characteristic length distribution; those with lengths <8 or >14 (for HLA-I) or >25 (for HLA-II) were filtered out. The resulting peptides were then checked for their source proteins. If a peptide came from a human protein, it was considered as a self peptide. If a peptide was identified by *de novo* sequencing and could not be found in the human protein database, i.e. a non-canonical HLA peptide, it was considered as a candidate neoantigen. The collection of self peptides was then used as the negative training data for that patient. For positive training data, we downloaded the table of T cell assays from IEDB with the following filters: linear epitopes and host as human. We further selected T cell epitopes that matched the patient’s alleles and had positive Assay Qualitative Measure, including Positive, Positive-High, Positive-Intermedia, Positive-Low. The resulting peptides were assumed to be recognized by the patient’s T cells, i.e. immunogenic, and were used to form the positive training data for that patient.

Using the above positive and negative training data, we then trained a personalized model, named DeepSelf, to predict whether a peptide is immunogenic or non-immunogenic. We used a bi-directional LSTM network coupled with amino acid embedding to model the peptides as they had been successfully used previously for de novo sequencing^27,52^, spectrum and retention time predictions^37,53,54^, and other peptide prediction tasks^8,55^. The model implementation was done in the TensorFlow^62^ framework (version 2.6). We used a Keras Sequential model that consists of an embedding layer of 8 neural units, a bi-directional LSTM layer of 8 units, a fully-connected layer with L2 regularizer, and a sigmoid activation layer. The model was trained using Adam optimizer and binary cross entropy loss for 100 epochs, only model weights with the best validation loss were saved. The Python implementation is available on GitHub (see the “Code availability” section).

The training dataset was splitted into three non-overlapping sets training-validation-testing with a ratio of 80-10-10. As the number of negative peptides was often several times higher than the number of positive peptides, we performed downsampling on negative peptides and trained 100 ensemble models per individual patient. The ensemble models were sorted according to their performance on the validation set and the average of the top-10 models was selected as the personalized prediction model for that patient. Finally, the model was applied to the candidate neoantigens, i.e. the non-canonical HLA peptides that could not be found in the human protein database. For each input peptide sequence, DeepSelf predicted a score from 0 to 1, where a higher score indicated that the peptide was more likely to be recognized by the patient’s T cells. The candidate neoantigens were sorted according to their DeepSelf scores and the top 10% were recommended for *in vitro* validations.

### Evaluation of DeepSelf and other immunogenicity prediction tools

We evaluated DeepSelf on the immunopeptidomes and neoantigens of 18 cancer patients from ten previously published studies^2,6,16,30–36^. The number of neoantigens varied from 2-6 per patient, all of which had been confirmed as immunogenic by T cell assay validations. The information of 18 patients and their datasets is summarized in Table S3; their neoantigens are provided in the Supplementary Data 3.

The key idea of DeepSelf is to use HLA self peptides obtained from MS-based immunopeptidomics as negative data to train immunogenicity prediction models. Four of the 18 patients, *Mel-15*, *Mel-0D5P*, *Mel-51*, and *iSPC2_IFN*, had MS-based immunopeptidomics data^6,30–32^. For each of those four patients, their data was analyzed using DeepNovo Peptidome and their HLA self peptides were obtained as described in the previous section. The number of HLA self peptides varied from 3,847-35,551 (Table S3). For the remaining 14 patients with no available MS data, we used their alleles to search the IEDB for peptides eluted by MHC assays in human. Those allele-matched MHC eluted peptides were then used instead of the unavailable self peptides as negative training data. Positive training data, i.e. T cell epitopes for each patient were obtained from the IEDB according to the patient’s alleles. The number of T cell epitopes varied from 133-4,785 (Table S3). The training datasets of 18 patients are provided in the Supplementary Data 4. Finally, personalized DeepSelf models were trained for each individual patient as described earlier; the trained models are provided on GitHub (see the “Code availability” section).

The personalized DeepSelf model of each individual patient was then evaluated on his/her own neoantigens. The number of neoantigens varied from 2-6 per patient, while the total number of peptides that had been tested in T cell assays was 15-434 per patient (Table S3). However, it should be noted that, before T cell assay validation, those peptides had actually been selected from an even larger pool of candidates^2,6,16,30–36^. Thus, we expanded the evaluation set of each patient by adding negative peptides randomly drawn from that patient’s self peptides (or allele-matched MHC eluted peptides from the IEDB if self peptides were not available). The ratio between neoantigens to negative peptides in the evaluation set of each patient was set at 1:100, i.e. an immunogenicity prediction tool needs to prioritize 1 neoantigen out of every 100 negative peptides. We found that the ratio 1:100 was in line with the number of neoantigens and the number of candidates identified from genomics/proteogenomics pipelines in most existing neoantigen studies^2,6,16,30–36^. Any common peptides shared between the evaluation set and the training data were excluded from the training data.

We compared DeepSelf to three other popular tools for immunogenicity prediction, including PRIME^23,24^ (version 2.0), NetMHCpan^4^ (version 4.1), and IEDB immunogenicity predictor^21^. Their prediction scores on the evaluation sets of 18 patients are provided in the Supplementary Data 5. Two criteria were used for the performance comparison. The first criterion is the area under the receiver operating characteristic curve (ROC-AUC) calculated from the prediction scores and the peptide classification in each evaluation set. The second criterion is the ranks of neoantigens versus negative peptides in each evaluation set based on the prediction scores.

The performance of four immunogenicity prediction tools are summarized in Figure 3. Figure 3a shows their prediction ROC curves and AUCs on the combined evaluation set of 18 patients. DeepSelf achieved an AUC of 0.69 and outperformed the other three tools. Figure 3b further shows the ability of the four prediction tools to rank neoantigens on top of non-immunogenic peptides. DeepSelf was able to rank 25% of the neoantigens within its top 10% predictions, and nearly 75% of the neoantigens within its top 30% (Q1 and Q3 quartiles of the boxplot, respectively). In other words, for an average patient evaluation set that included 4 neoantigens and 400 non-immunogenic peptides, DeepSelf’s top 40-120 predicted candidates would contain 1-3 neoantigens of interest. Figure 3b also shows that DeepSelf again outperformed the other tools in terms of the ranking ability.

Figure 3c further shows the prediction tools’ performance on each individual patient. DeepSelf outperformed the other tools on 12 out of 18 patients. One of the best prediction results of DeepSelf was achieved on *Mel-15* (AUC=0.85), a melanoma patient dataset first published by Bassani-Sternberg et al.^6^. This dataset is well-known as the native immunopeptidomics sample with the highest coverage to date, where more than 35K self peptides had been identified by MS-based immunopeptidomics. Bassani-Sternberg et al.^6^ and Wilhelm et al.^37^ identified 28 mutated HLA-I peptides from the MS and RNA-seq data of *Mel-15*, and among them, five neoantigens were confirmed as immunogenic by T cell assay validation. Figure 3d shows the predicted ranks of those five neoantigens versus other non-immunogenic peptides identified from that patient. DeepSelf produced the best ranks across all five neoantigens, and moreover, three of them were ranked within its top 3% predictions.

### *In vitro* validation of autologous neoantigen-specific T cell responses

IFN-γ ELISPOT was conducted to validate the responsiveness of putative neoantigens to CD8+ T cells. Cryopreserved Peripheral blood mononuclear cells of patient were recovered for IFN-γ ELISPOT, and every reaction of antigen stimulation was performed three times. Total of 100 neoantigens were synthesized (purity>95%). 20 neoantigen-mix were used(2μg/ml) in the first screening. Then the cells from positive ELISPOT wells were expanded up to 80% with 200 IU recombined human IL-2 and ELISPOT-positive peptide mix(2μg/ml). ELISPOT screening were further performed on expanded cells for 4 neoantigen mix and then single neoantigen. Finally, flow cytometry and ELISPOT were performed three times for each reactive neoantigen to detect the 4-1BB expression and confirm the reaction again, respectively.

In T cell validation experiments, the CD137, IFN-γ and ELISPOT were conducted with 3 replicates. The experiment groups were compared to the control. The P-value was calculated using a two-sided unpaired Student-t test. All the analysis was performed with GraphPad Prism Version 8.2.1.

## Data availability

The HLA Ligand Atlas dataset was obtained from Marcu et al.^29^ and the ProteomeXchange Consortium via the PRIDE^63^ partner repository with the dataset identifier PXD019643. The mass spectrometry proteomics data of five cervical cancer patients generated in our study have been deposited to the ProteomeXchange Consortium via the PRIDE^63^ partner repository with the dataset identifier PXD046370.

## Software and code availability

The Python source code of DeepSelf is publicly available at https://github.com/nh2tran/DeepImmun. DeepNovo Peptidome is available in PEAKS Studio and PEAKS Online at https://www.bioinfor.com/peaks-software/.

## Acknowledgements

This work was partially supported by the following grants: National Key R&D Program of China, grant No. 2022YFA1304603 (C.P.), Canada NSERC OGP0046506 (M.L.), the Canada Research Chair program (M.L.), and the NSFC No. 61832019 (M.L.). Y.Z. was supported by the National Key R&D Program: Intergovernmental International Science and Technology Innovation Cooperation Project (Grant No. 2022YFE0141000),the National Natural Science Foundation of China (Grant no. 82272873), Medical Science and Technology Project of Henan Province (Grant no. SBGJ202101010), Project of Central Leading Local Science and Technology Development of Henan Province (Grant no. Z20221343036).

## Author contributions

Ngoc Hieu Tran designed and trained DeepNovo Peptidome and DeepSelf models.

Chao Peng designed the immunoprecipitation and LC-MS/MS experiments, and Haofei Miao and Ping Wu performed the experiments.

Haiming Qin and Jingxiang Lang prepared the patient samples and performed *in vitro* validation of T cell responses.

Lei Xin, Rui Qiao, Qing Zhang, Wenting Li, Ping Wu, Chao Peng and Baozhen Shan developed and tested DeepNovo Peptidome.

Qingyang Lei, Dongbo Bu, Haicang Zhang, Chungong Yu, and Xiaolong Liu contributed to the sample preparation, data analysis, and result validation.

Ngoc Hieu Tran, Chao Peng, Haiming Qin, Qingyang Lei, Baozhen Shan, and Ming Li wrote the manuscript.

Ngoc Hieu Tran, Baozhen Shan, and Ming Li proposed and planned the study. Yi Zhang, Baozhen Shan, and Ming Li supervised the study.

---

**Fig. S1.**
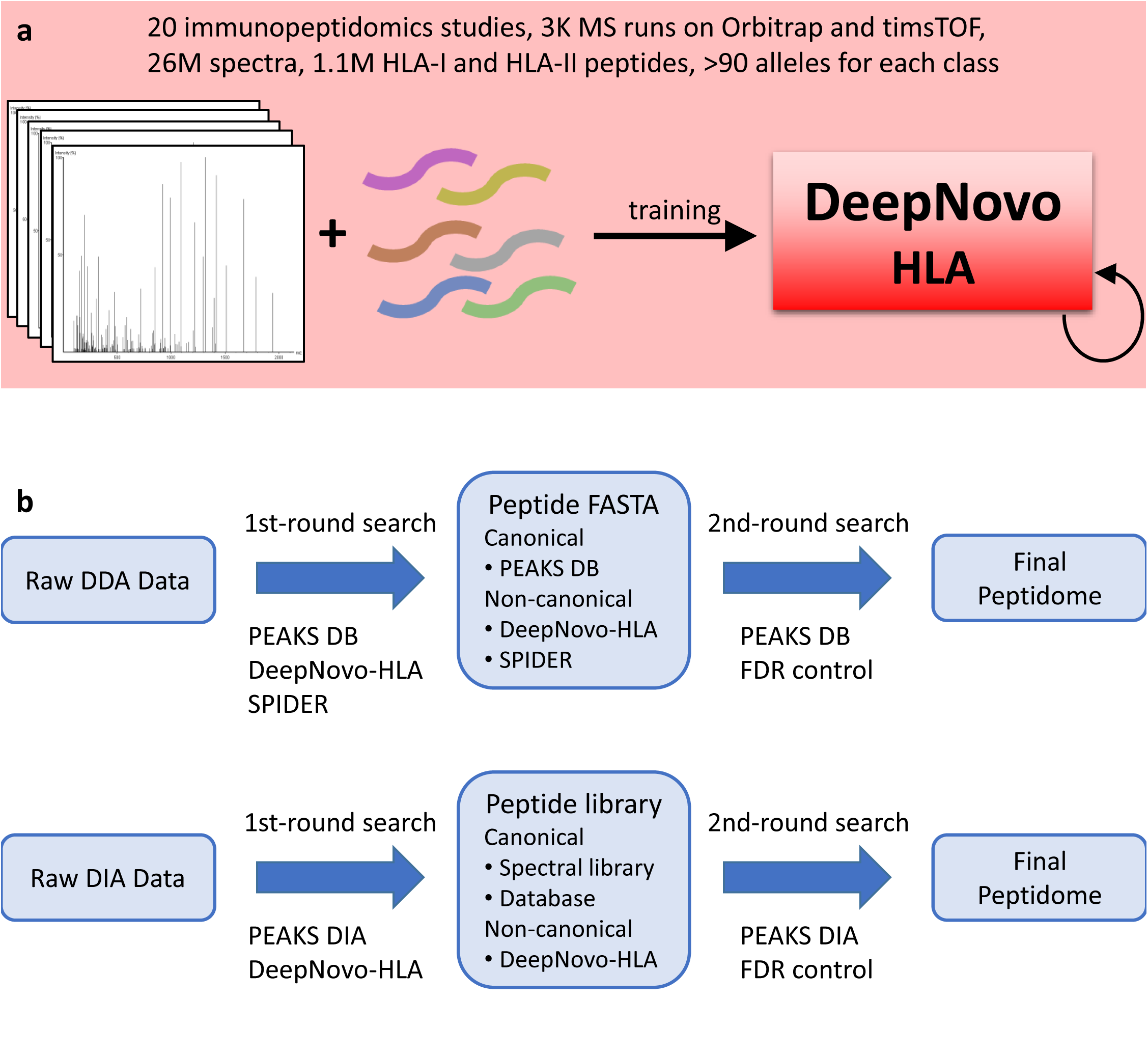
DeepNovo Peptidome workflow. **a**, DeepNovo-HLA: a specialized model for de novo sequencing of HLA peptides. **b**, DeepNovo Peptidome workflow for DDA- and DIA-MS immunopeptidomics. (MS: Mass Spectrometry; HLA: Human Leukocyte Antigen; DDA: Data-dependent Acquisition; DIA: Data-independent Acquisition; FDR: False Discovery Rate)

