## Supplementary Data 1 for "Discovering and Validating Neoantigens by Mass Spectrometry-based Immunopeptidomics and Deep Learning"

Supplementary Data 1 can be downloaded from the temporary link below. 
It is too big (135 MB) to upload to the journal website for an initial submission.
We shall work with the journal for a permanent link.

Download link:
https://drive.google.com/file/d/1o_5AVk1Qvr-qfrXjAz09BUDapTDLY3VW/view?usp=sharing
