## Supplementary Data 4 for "Discovering and Validating Neoantigens by Mass Spectrometry-based Immunopeptidomics and Deep Learning"

Supplementary Data 4 can be downloaded from the temporary link below. 
It is too big to upload to the journal website for an initial submission.
We shall work with the journal for a permanent link.

Download link:
https://drive.google.com/file/d/1FCCTZoSuujIOe1U7B-dpuUpVnPI93-hX/view?usp=sharing
